# Sequential action of the nuclear Argonautes HRDE-1 and NRDE-3 promotes global transcriptional silencing during starvation in the nascent *C. elegans* germline

**DOI:** 10.1101/2025.06.25.661390

**Authors:** Emilie Chien, Mezmur D. Belew, W. Matthew Michael

## Abstract

In *C. elegans*, if embryos hatch into an environment lacking nutrients the primordial germ cells (PGCs) will arrest mRNA transcription and enter quiescence until food becomes available. Our previous work had shown that the global transcriptional silencing (GTS) that occurs in PGCs during L1 starvation requires a “hyperdeposition” of the repressive histone mark H3K9me3 in germline chromatin. Hyperdeposition is defined as an increase in the amount of H3K9me3 embedded in germline chromatin, relative to neighboring somatic nuclei. Hyperdeposition initiates in embryonic PGCs and is maintained in starved L1s. Here, we show that the nuclear RNAi pathway contributes to both the initiation and maintenance of H3K9me3 hyperdeposition. Interestingly, both known nuclear Argonautes, HRDE-1 and NRDE-3, play a role in hyperdeposition and they do so in a sequential manner. We show that HRDE-1 acts in embryonic PGCs to initiate hyperdeposition and then, at the embryo-to-L1 transition, HRDE-1 becomes dispensable and NRDE-3 is required to maintain H3K9me3 levels, and for GTS. We also examine the timing of H3K9me3 hyperdeposition and find that it initiates soon after germline zygotic genome activation (ZGA) occurs. Our data suggest a model where ZGA promotes both gene expression and H3K9me3 deposition at active loci, and under starvation conditions these H3K9me3 marks are then employed to silence the germline genome.

## Introduction

Gene expression controls cell function and fate. How gene expression is regulated has been the topic of numerous studies over the past several decades, and most have examined gene expression on a small scale, at the level of individual genes or multi-genic operons. However, much less is known about how mRNA transcription is controlled at the genome scale.

The nematode *Caenorhabditis elegans* is a premier model system to study the global regulation of gene expression. Recent work from our group has shown that the *C. elegans* germline undergoes multiple cycles of global transcriptional activation and silencing during development (Belew et al., 2021; Belew & Chien et al., 2023; Chien & Michael, 2023; Belew & Chien et al., 2024). The germline precursor cells, also known as the P-lineage, are globally silenced by the OMA-1/2 proteins at the one– to two-cell stages, and by PIE-1 for the remainder of the P-lineage divisions (Wang & Seydoux, 2013). P_4_, the last of the P-lineage cells, divides to create the primordial germ cells (PGCs) Z2 and Z3 (Z2/Z3). Z2/Z3 are transcriptionally active but arrest the cell cycle shortly after their birth and remain arrested through the remainder of embryogenesis into the first L1 larval stage (Fukuyama et al., 2003).

*C. elegans* are able to developmentally arrest growth, a condition known as diapause, when facing adverse environmental conditions such as the lack of nutrients. One such state occurs during the L1 stage, called L1 diapause, and is distinct from the well-studied dauer stage (Baugh & Hu, 2020). Animals in L1 diapause, or “starved L1s”, will remain arrested until food becomes available, and are able to survive for weeks in this condition. Our group has previously shown that one manifestation of L1 diapause is an event we refer to as global transcriptional silencing (GTS), where the energy sensing kinase AMPK directs a global chromatin compaction pathway to compact germline chromatin and to extinguish RNA polymerase II (RNAPII) mediated transcription (Belew et al., 2021). Components of this compaction pathway include the TOP-2/condensin II axis, which promotes loop extrusion, as well as members of the H3K9me3/heterochromatin pathway (Belew et al., 2021). Indeed, when the repressive H3K9me3 chromatin mark is examined in starved L1s, we observed a hyperdeposition of this mark on Z2/Z3 chromatin, relative to neighboring somatic cells (Belew et al., 2021). These published data showed that H3K9me3 hyperdeposition is a key event leading to Z2/Z3 GTS, and the focus of the current study was to define how this occurs.

Nuclear RNA interference (RNAi) pathways are known to regulate gene expression through a variety of processes including silencing transposons, facilitating epigenetic inheritance, and acting via both co-transcriptional and post-transcriptional silencing mechanisms (Castel & Martienssen, 2013; Youngman & Claycomb, 2014). The small RNA (sRNA)-guided NRDE pathway is of particular interest as this pathway can shut down gene expression in a targeted manner. More specifically, in the NRDE pathway, an sRNA-bound nuclear Argonaute protein recruits downstream factors NRDE-1, NRDE-2, and NRDE-4 for targeted deposition of H3K9me3 (Castel & Martienssen, 2013; Youngman & Claycomb, 2014). Thus, the NRDE pathway couples active transcription by RNAPII to the silencing effect of H3K9me3. Our work has demonstrated that embryonic Z2/Z3 are actively transcribing, while also undergoing hyperdeposition of the repressive H3K9me3 mark (Belew et al., 2021). Are genes that are transcribed in embryonic Z2/Z3 the targets for H3K9me3 hyperdeposition? The co-transcriptionally acting NRDE pathway may hold the answer to this question.

In this study, we explore a novel role of the nuclear RNAi NRDE pathway in silencing transcription in a genome-wide manner in *C. elegans* primordial germ cells. We employ genetics and cytology to examine Z2/Z3 transcriptional behavior and to identify the role of NRDE pathway components in Z2/Z3 during embryogenesis and L1 larval stage. Altogether, our findings demonstrate that the NRDE pathway facilitate the global deposition of H3K9me3 in Z2/Z3, and that different nuclear Argonautes direct the pathway in a life stage-dependent manner.

## Results

### PIE-1 is degraded and transcription is activated immediately upon the birth of Z2/Z3

In *C. elegans*, germline precursor cells (i.e. the P lineage) are transcriptionally repressed by the zinc-finger protein PIE-1 (Seydoux et al., 1996; Mello et al., 1996). PIE-1 is thought to inhibit mRNA transcription by blocking the C-terminal repeat domain phosphorylation on RNAPII, thereby preventing elongation (Seydoux & Dunn, 1997). Previous work showed that, upon the division of P_4_ to form the Z2/Z3 PGCs, PIE-1 is abruptly degraded and Z2/Z3 become transcriptionally active (Mello et al., 1996; Wang & Seydoux, 2013). These findings suggest a simple model whereby PIE-1 destruction is the trigger for zygotic genome activation (ZGA) in the nascent germline. However, the precise timing of PIE-1 degradation and RNAPII activation has not been delineated. To study this, we developed a framework to stage Z2/Z3 during gastrulation in the early embryo. We used immunofluorescence (IF) staining with an antibody against the germ cell specific P-granules to monitor Z2/Z3 from their birth through the end of gastrulation (Fig S1). At the 88-cell stage, P_4_ divides to form Z2/Z3, and these cells are born at the ventral surface and are positioned parallel to the anterior-posterior axis. We designated this as the <u>Pre-I</u>ngression (Pre-I) developmental stage (Fig S1). Next, as gastrulation commences, Z2/Z3 ingress into the center of the embryo, while remaining parallel to the anterior-posterior axis. We named this the <u>Post-</u>Ingression, <u>Pre-R</u>eorientation (Post-I/Pre-R) stage (Fig S1). Finally, at the end of gastrulation, Z2/Z3 reorient to become perpendicular to the anterior-posterior axis, and we named this the <u>Post-I</u>ngression, <u>Post-R</u>eorientation (Post-I/Post-R) embryonic stage (Fig S1).

With the early embryonic Z2/Z3 stages established, we next asked: when is PIE-1 degraded and when does transcription begin in Z2/Z3? Using a *gfp::pie-1* strain where *gfp* is integrated at the endogenous *pie-1* locus (Kim et al., 2014), we stained embryos for GFP to monitor PIE-1. We note that this GFP::PIE-1 protein is fully functional (Kim et al., 2021). To monitor RNAPII activity we co-stained for the presence of phospho-serine 2 within RNAPII’s CTD (RNAPIIpSer2), as this mark correlates with active and elongating RNAPII and has been used extensively in the field to track RNAPII activity *in situ* (Palancade & Bensaude, 2003; Larson et al., 2016; Belew et al., 2021). As shown in Fig 1A, P_4_ has strong GFP::PIE-1 presence, and no detectable RNAPIIpSer2, indicating that P_4_ is not transcriptionally active, as expected. Interestingly, we see that at the Pre-I stage, both GFP::PIE-1 and RNAPIIpSer2 signals are detectable in Z2/Z3, in a non-mutually exclusive manner (Fig 1B, 1F). By the Post-I/Pre-R stage, however, GFP::PIE-1 has all but disappeared from Z2/Z3, and Z2/Z3 are now actively transcribing (Fig 1C-D). We see a significant decrease in the signal intensity of GFP::PIE-1 from the P_4_ to Pre-I stage, as well as from the Pre-I to Post-I/Pre-R stage (Fig 1G). There is a corresponding significant increase in the signal intensity of RNAPIIpSer2 in Z2/Z3 from the P_4_ to Pre-I stage (Fig 1H). We conclude that GFP::PIE-1 disappears from, and RNAPIIpSer2 appears in, Z2/Z3 shortly after they are born at the Pre-I stage, and that the absence or presence of each signal is complete by the time Z2/Z3 have ingressed at the Post-I/Pre-R stage. In sum, this data suggests that PIE-1 is degraded and that transcriptional activation begins immediately after the birth of Z2/Z3, during the Pre-I stage. Similar results have been reported by others (Schaner et al., 2003; Checchi & Kelly, 2006).

**Figure 1:**
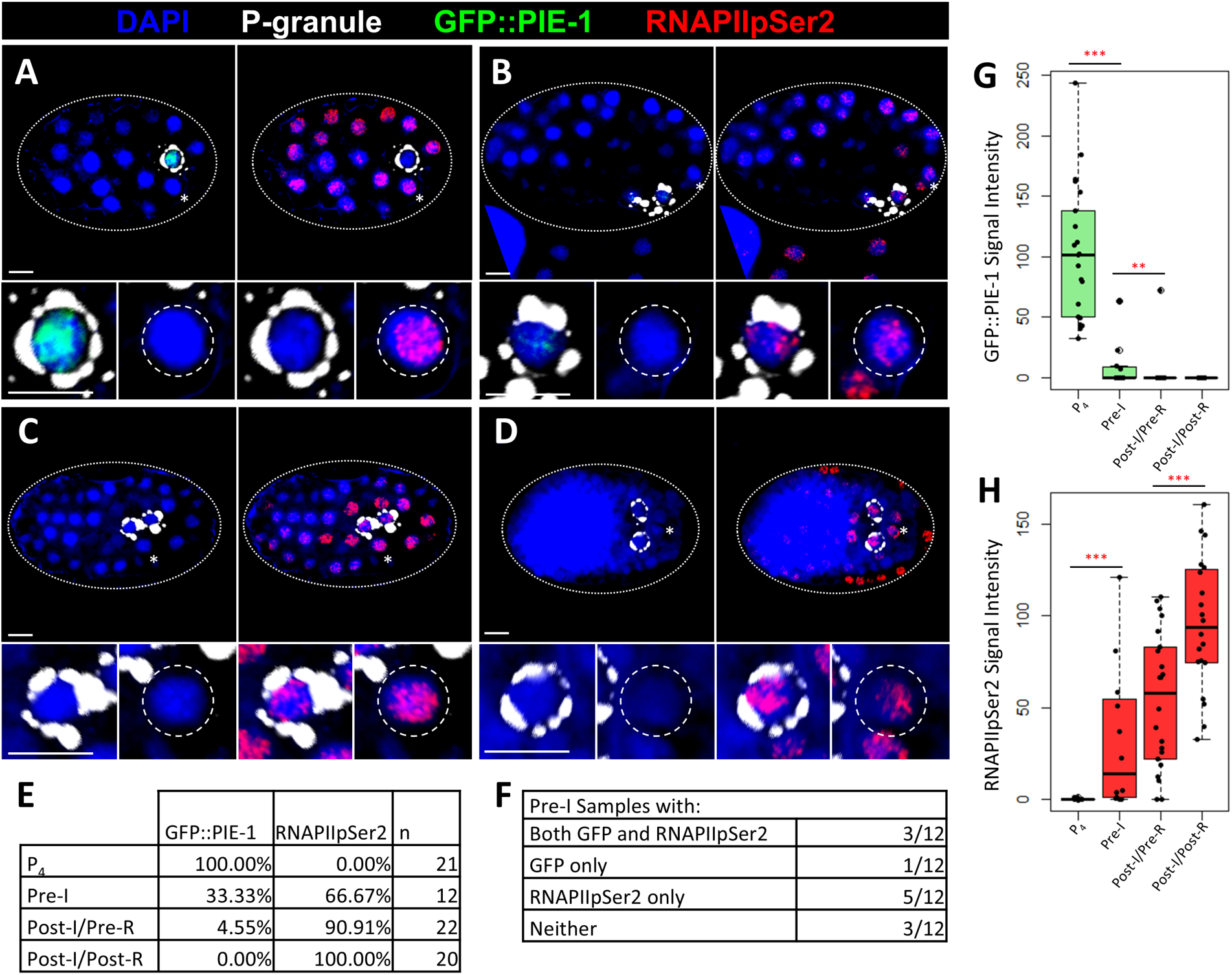
Activation of transcription occurs upon destruction of PIE-1 in Z2/Z3. **(A-D)** GFP::PIE-1 embryos were co-stained for DAPI (blue), P-granules (white), GFP (green), and RNAPIIpSer2 (red). Insets show P_4_ or Z2/Z3 (left, identified by P-granules) and a nearby somatic cell (right). The four embryonic stages shown are (A) P_4_, (B) Pre-I, (C) Post-I/Pre-R, and (D) Post-I/Post-R. Dotted lines indicate embryo outline, dashed lines indicate somatic cell outline. Scale bars: 5 μm. **(E)** P_4_ and Z2/Z3 were scored for the presence of GFP::PIE-1 and RNAPIIpSer2 signal at various embryonic stages. **(F)** Z2/Z3 in Pre-I embryos were scored for the independent or combined presence of GFP and RNAPIIpSer2. **(G-H)** Quantification of normalized raw integrated density of GFP and RNAPIIpSer2 for data presented in A-F. *p≤0.05; **p≤0.01; ***p≤0.001

### Initial onset of transcription in Z2/Z3 is regulated by *zif-1* dependent degradation of PIE-1

Now that we have determined the timing of PIE-1 degradation and initial onset of transcription, we next asked if transcription initiation is dependent on the degradation of PIE-1. PIE-1 degradation in Z2/Z3 has been shown to be dependent on zinc-finger interacting protein ZIF-1 (Reese et al., 2000, Checchi & Kelly 2006). ZIF-1 is an E3 ubiquitin ligase substrate recruitment factor and ubiquitinates PIE-1 for degradation (DeRenzo et al., 2003). Using *zif-1* (RNAi) or *zif-1^-/-^* embryos, we co-stained for GFP::PIE-1 and RNAPIIpSer2, with a focus on the Pre-I stage (Fig 2A-C). Here, we expect that if the initial activation of transcription is dependent on PIE-1 degradation, then the appearance of the RNAPIIpSer2 signal would be delayed in *zif-1* (RNAi) or *zif-1^-/-^* embryos. Indeed, we found that under both conditions, *zif-1* (RNAi) or *zif-1^-/-^*mutants, there was a significant increase in detectable GFP::PIE-1, and concomitant decrease in the RNAPIIpSer2 signal (Fig 2A-C). We note that the timing of PIE-1 degradation after loss of ZIF-1 is only modestly delayed, as reported by others (Checchi & Kelly, 2006), showing that Z2/Z3 contain a redundantly acting PIE-1 destruction pathway. Nonetheless, the minor delay in ZIF-1 destruction is sufficient to repress RNAPII activation, showing that PIE-1 destruction is indeed rate-limiting for the activation of RNAPII in the Z2/Z3 PGCs.

**Figure 2:**
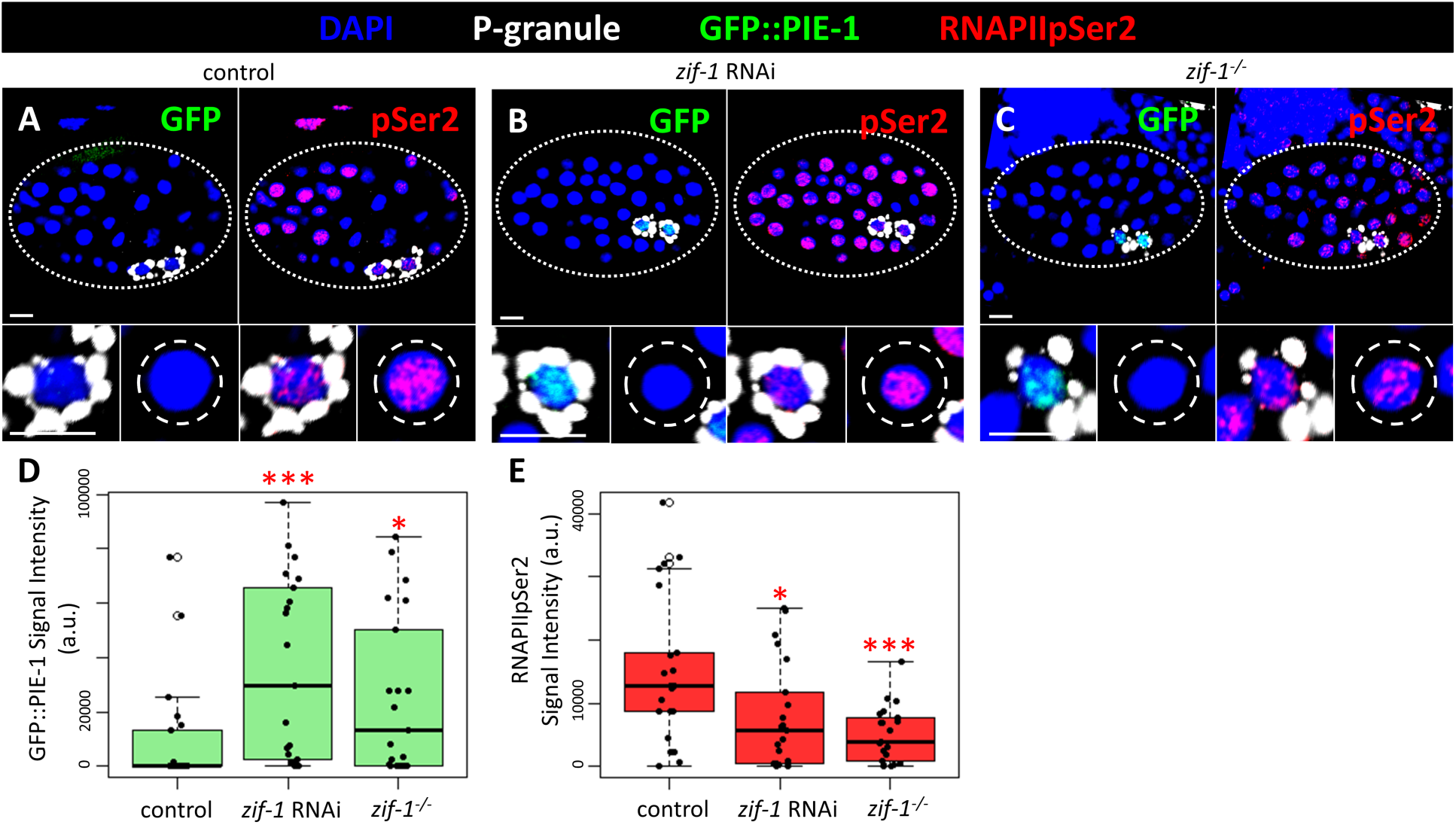
Depletion or absence of ZIF-1 delays PIE-1 destruction and initial activation of transcription in Z2/Z3. **(A-C)** GFP::PIE-1 Pre-I stage embryos were co-stained for DAPI (blue), P-granules (white), GFP (green), and RNAPIIpSer2 (red). Insets show P_4_ or Z2/Z3 (left, identified by P-granules) and a nearby somatic cell (right). The three conditions shown are (A) control, (B) *zif-1* RNAi, (C) and *zif-1^-/-^*. Dotted lines indicate embryo outline, dashed lines indicate somatic cell outline. Scale bars: 5 μm. **(D-E)** Quantification of normalized raw integrated density for data presented in A-C. n=21; *p≤0.05; **p≤0.01; ***p≤0.001

### H3K9me3 hyperdeposition in Z2/Z3 initiates soon after the onset of gastrulation

In previous work, we found that Z2/Z3 chromatin accumulate H3K9me3 to a much greater extent than do surrounding somatic nuclei (Belew et al., 2021). We refer to this phenomenon as “H3K9me3 hyperdeposition”. Our previous work showed that H3K9me3 hyperdeposition occurs sometime during embryogenesis, although the precise timing was not determined. To examine this, and to correlate hyperdeposition to germline ZGA, we stained wild-type embryos for H3K9me3 and focused on the P_4_ through the Post-I/Post-R stages. As shown in Fig 3A, in P_4_ H3K9me3 manifests as small patches scattered throughout the DAPI-stained DNA. Just after Z2/Z3 are born (the Pre-I stage), the pattern is similar, however by the time gastrulation begins (Post-I/Pre-R) the signal increases such that more of the DAPI-stained DNA contains H3K9me3 signal (Fig 3A). By the time gastrulation is complete (Post-I/Post-R) H3K9me3 hyperdeposition is also complete. To quantify this, we simply measured the area of the H3K9me3 signal in single, maximum-intensity confocal images and divided this by the area of the DNA-based DAPI signal. As shown in Fig 3B, the amount of DNA covered by H3K9me3 signal gradually increased from the birth of Z2/Z3 through gastrulation. Fig 3C summarizes our findings thus far; we found that at the P_4_ stage PIE-1 is present, RNAPII is silent, and H3K9me3 has yet to be hyperdeposited. Upon the birth of Z2/Z3, PIE-1 is abruptly degraded, RNAPII is activated, and H3K9me3 hyperdeposition initiates. By the end of gastrulation RNAPII is active and H3K9me3 hyperdeposition is complete. These data thus provide a temporal correlation between the onset of RNAPII transcription and H3K9me3 hyperdeposition in the Z2/Z3 PGCs.

**Figure 3:**
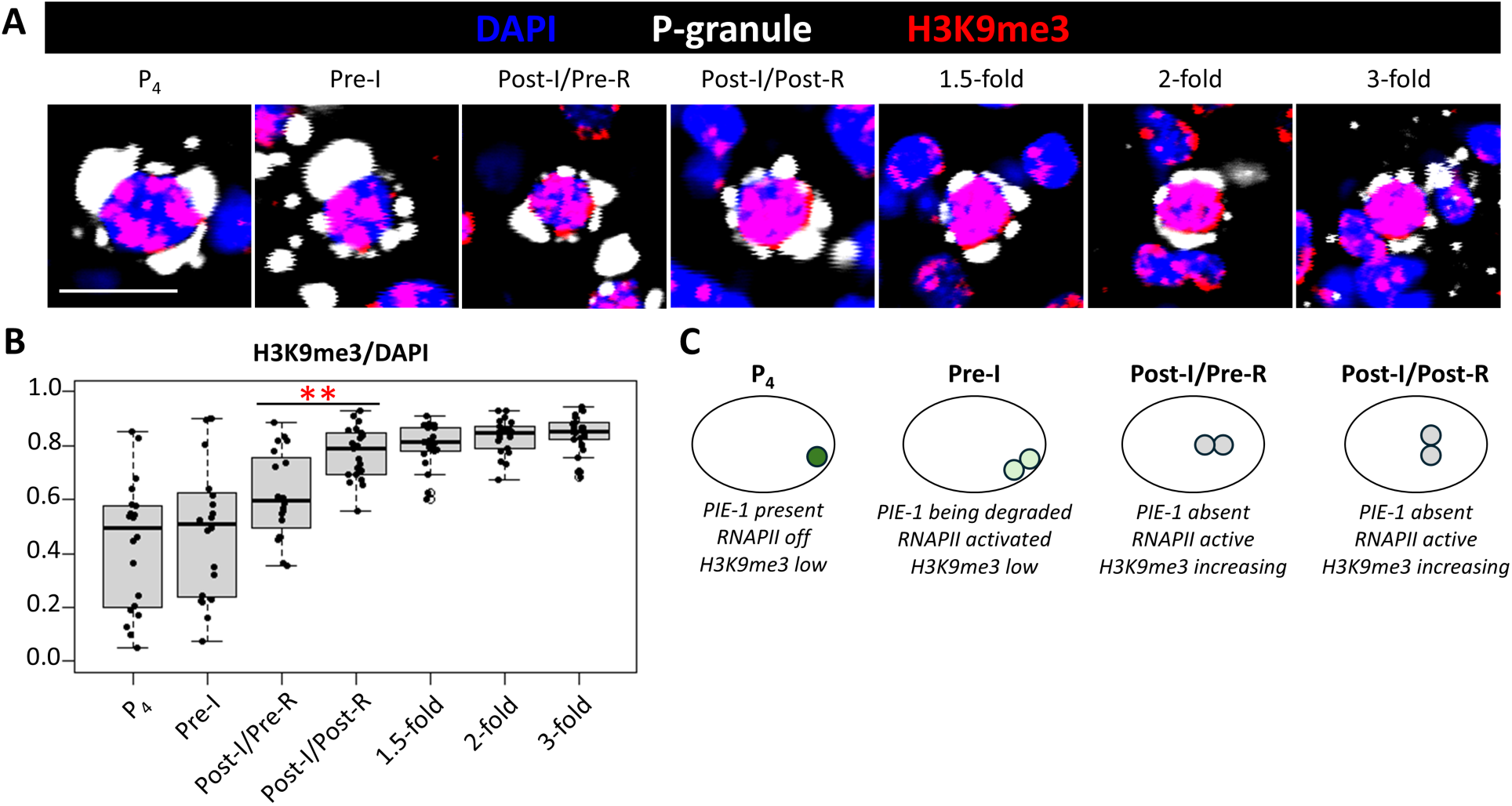
Hyperdeposition of H3K9me3 in Z2/Z3 is complete by Post-I/Post-R stage. **(A-D)** Wild-type embryos were stained for DAPI (blue), P-granules (white), and H3K9me3 (red). The four embryonic stages shown are (A) P_4_, (B) Pre-I, (C) Post-I/Pre-R, and (D) Post-I/Post-R. Scale bar: 5 μm. **(B)** Quantification of H3K9me3 signal overlap with DAPI in P_4_ or Z2/Z3 for data presented in A-D. n=20; *p≤0.05; **p≤0.01; ***p≤0.001 **(C)** Summary of PIE-1, RNAPII, and H3K9me3 signal at various embryonic stages.

### H3K9me3 hyperdeposition in Z2/Z3 is dependent on the HRDE-1 guided NRDE pathway during embryogenesis

We next investigated the molecular machinery responsible for Z2/Z3 H3K9me hyperdeposition and considered the sRNA-guided NRDE pathway. In this pathway, the Argonautes (AGOs) HRDE-1 or NRDE-3 uses sRNA as guides to identify target loci that are undergoing active transcription. Base-pairing between the sRNA and a nascent mRNA allows recruitment of HRDE-1/NRDE-3 to the locus, where the remainder of the NRDE pathway (NRDE-1, –2, and –4) assembles to promote H3K9me3 deposition at nearby nucleosomes. Because H3K9me3 hyperdeposition is coincident with RNAPII transcription in Z2/Z3, the NRDE pathway is a good candidate to promote hyperdeposition in this system. To examine this, we stained Post-I/Post-R stage Z2/Z3 for H3K9me3 and DNA in wild type and *nrde* mutant embryos. Because the efficiency of IF staining can vary from one slide to the next, we mounted both wild type and mutant samples on the same slide, thereby ensuring that IF efficiency was identical across both sample sets. Our wild type strain expresses GFP-tagged PGL-1, allowing us to distinguish wild type from mutant strains. We measured the areas of both H3K9me3 and DNA signals and computed their ratios, as in Fig 3. As shown in Fig 4A, *nrde-4* mutants were significantly decreased for H3K9me3 deposition, relative to wild type. This was also true for *hrde-1* mutants (Fig 4B), but not for a strain harboring the HK->AA mutations in *nrde-3* (Fig 4C). These mutations have been shown previously to disable sRNA binding by NRDE-3 (Chen and Phillips, 2025). We also examined *set-25* mutants and found that H3K9me3 was nearly undetectable, as expected (Fig 4D). Taken together, these data show that some H3K9me3 deposition in the Z2/Z3 PGCs is promoted by the NRDE pathway, and that HRDE-1 and not NRDE-3 is the relevant secondary AGO.

**Figure 4:**
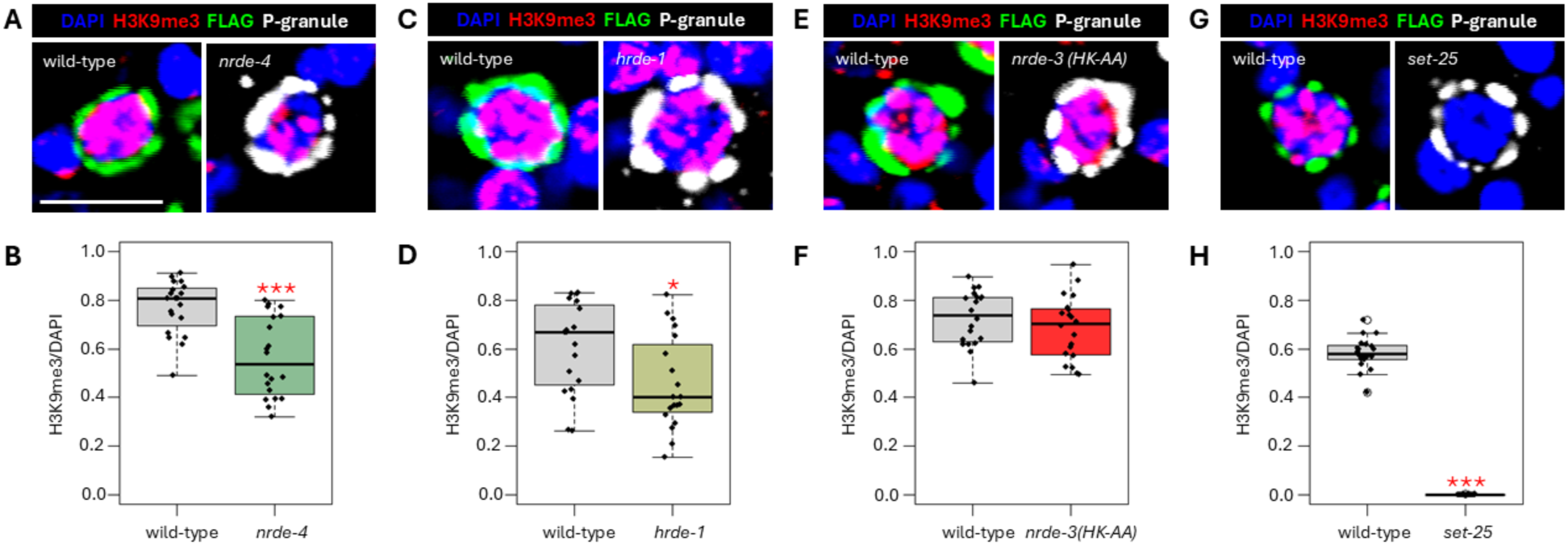
HRDE-1 is required for hyperdeposition of H3K9me3 in Z2/Z3 in embryos. **(A-E)** Post-I/Post-R embryos were stained for DAPI (blue), P-granules (white), and H3K9me3 (red) in wild-type, *nrde-3 (HK-AA)*, *hrde-1*, *nrde-4*, and *set-25* mutant strains. Scale bar: 5 μm. **(F)** Quantification of H3K9me3 signal overlap with DAPI in Z2/Z3 for data presented in A-E. *p≤0.05; **p≤0.01; ***p≤0.001

### The NRDE pathway is required for maintenance of H3K9me3 hyperdeposition in Z2/Z3 of starved L1s

Our previous work had shown that H3K9me3 levels on Z2/Z3 chromatin remain high, relative to the soma, after embryos hatch and they remain high if L1s undergo starvation. Furthermore, our previous work showed that H3K9me3 marks are required for GTS during L1 starvation (Belew et al., 2021). It was, therefore, important to determine if the NRDE pathway was also active in Z2/Z3 during L1 starvation. We starved *nrde-1^-/-^*, *nrde-2^-/-^*, and *nrde-4^-/-^* mutant L1s and then stained Z2/Z3 for H3K9me3 and DNA. As in Fig 4, wild type and mutant samples were mounted on the same slide and wild type was identified via the presence of GFP-tagged PGL-1. We observed significantly reduced levels of H3K9me3 signal, relative to wild type, in *nrde-1* and *nrde-2* mutants (Figs 5A-D), but not in *nrde-4* or *hrde-1* mutants (Figs 5E-H). As expected, in *set-*25 mutants the H3K9me3 signal was nearly undetectable (Figs I&J). We also examined a double mutant for *ampk-1* and *ampk-*2, as our previous work had shown that these two genes are required for GTS during L1 starvation (Belew et al, 2021), and found H3K9me3 levels in the double mutant were significantly higher than wild type (Figs 5K-L). Taken together, these finding show that some *nrde* pathway components, like *nrde-1* and *nrde-2*, are required for the maintenance of H3K9me3 levels during L1 starvation, whereas other, like *hrde-1* and *nrde-4*, are not. In addition, the data show that AMPK-1/2 act downstream of H3K9me3 deposition to promote GTS during L1 starvation.

**Figure 5:**
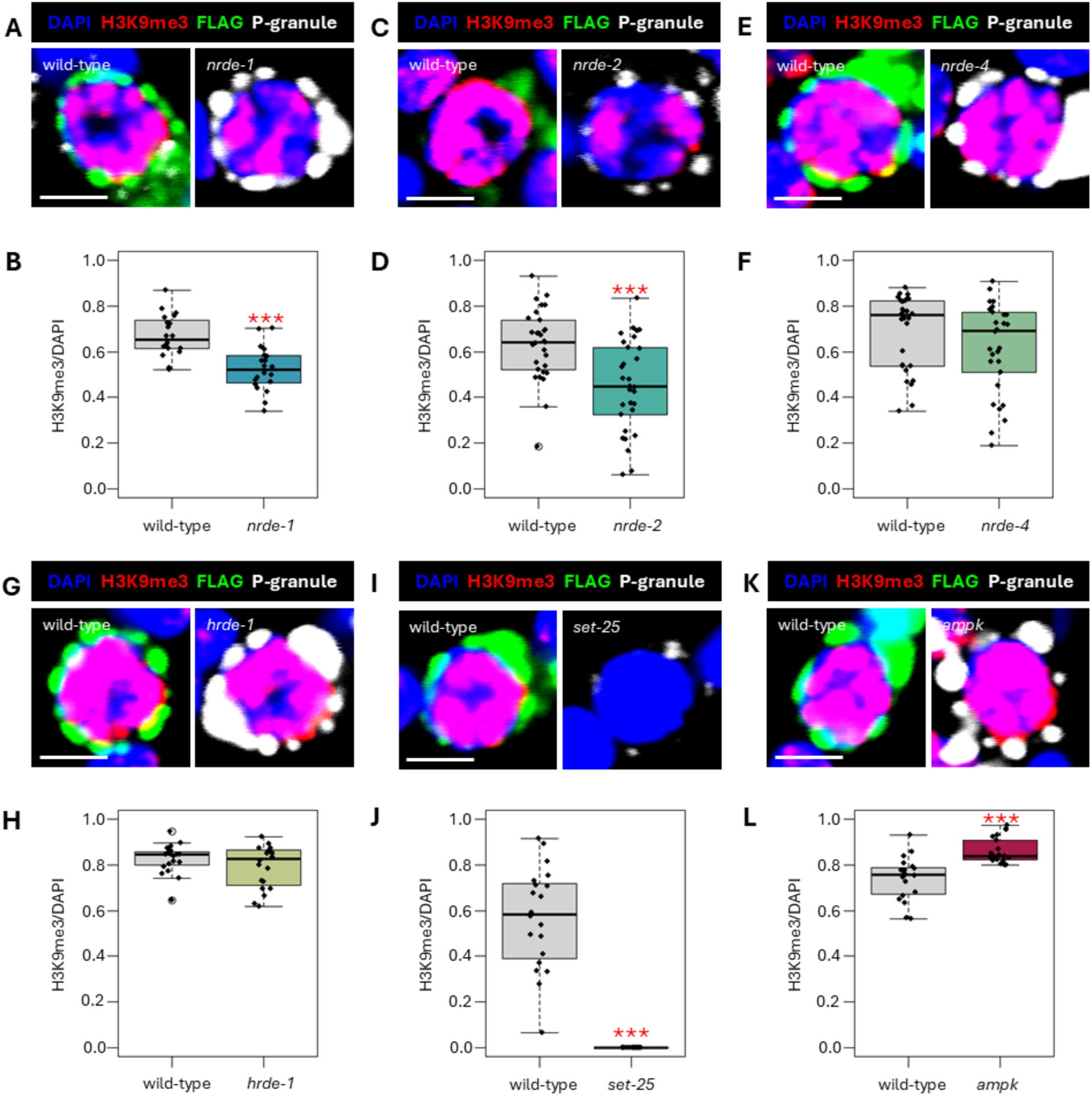
NRDE pathway components, excluding argonaute HRDE-1, are required for H3K9me3 hyperdeposition in Z2/Z3 in starved L1s. **(A)** Z2/Z3 of starved L1s were stained for DAPI (blue), P-granules (white), and H3K9me3 (red) in wild-type, *nrde-1*, *nrde-2*, *nrde-4*, *hrde-1, hpl-2 (RNAi), nrde-1 + hpl-2 (RNAi),* and *set-25* strains. Scale bar: 5 μm. **(B)** Quantification of H3K9me3 signal overlap with DAPI for Z2/Z3 of starved L1s for data presented in (A). *p≤0.05; **p≤0.01; ***p≤0.001

### NRDE-3, not HRDE-1, is responsible for maintaining H3K9me3 in Z2/Z3 in starved L1s

Of *C. elegans* nuclear Argonautes, only two are known to work within the NRDE pathway – HRDE-1 and NRDE-3 (Seroussi et al., 2023). Since HRDE-1 is dispensable for H3K9me3 hyperdeposition in Z2/Z3 in starved L1s, we next asked if NRDE-3 assumes this role. Using the NRDE-3 sRNA-binding mutant *nrde-3 (HK-AA),* we stained starved L1 animals for H3K9me3. Interestingly, we observed that *nrde-3 (HK-AA)* animals had significantly less H3K9me3 signal, relative to wild-type, indicating that NRDE-3 is the Argonaute responsible for H3K9me3 hyperdeposition in Z2/Z3 of starved L1s (Fig 6A&B). In sum, our data show that HRDE-1 is the Argonaute for H3K9me3 hyperdeposition in embryonic Z2/Z3, and that there is a hand-off at the embryo-to-L1 transition to NRDE-3 to maintain the marks. To explore this further we examined NRDE-3 expression in embryos and starved L1s, using a strain where *gfp* had been integrated into the endogenous *nrde-3* locus. We note that this GFP::NRDE-3 fusion protein retains all function (Seroussi et al., 2023). As shown in Fig S2A-B, GFP::NRDE-3 is expressed at low levels in P_4_, and expression appears to decline such that signal is difficult to detect in gastrulating Z2/Z3. GFP signal recovers somewhat after gastrulation, and by the time embryos have hatched and L1s starve GFP signal is easily detectable in Z2/Z3. It thus appears that NRDE-3 levels are reduced in embryonic Z2/Z3 and then come back during L1 starvation, perhaps explaining the differential requirements for NRDE-3 function in embryos and starved L1s. We also examined HRDE-1 localization in embryos and found that expression is limited to PGCs where it is present and localized to nuclei in all conditions examined (Fig S2C). These data show that embryonic PGCs express HRDE-1 during H3K9me3 hyper-deposition, but not NRDE-3, and this is consistent with our data showing the *hrde-1* and not *nrde-3* is required for the hyper-deposition (Fig 4). During L1 starvation a transition occurs, where despite both NRDE-3 and HRDE-1 being present in PGCs only *nrde-3* is required for hyper-deposition.

**Figure 6:**
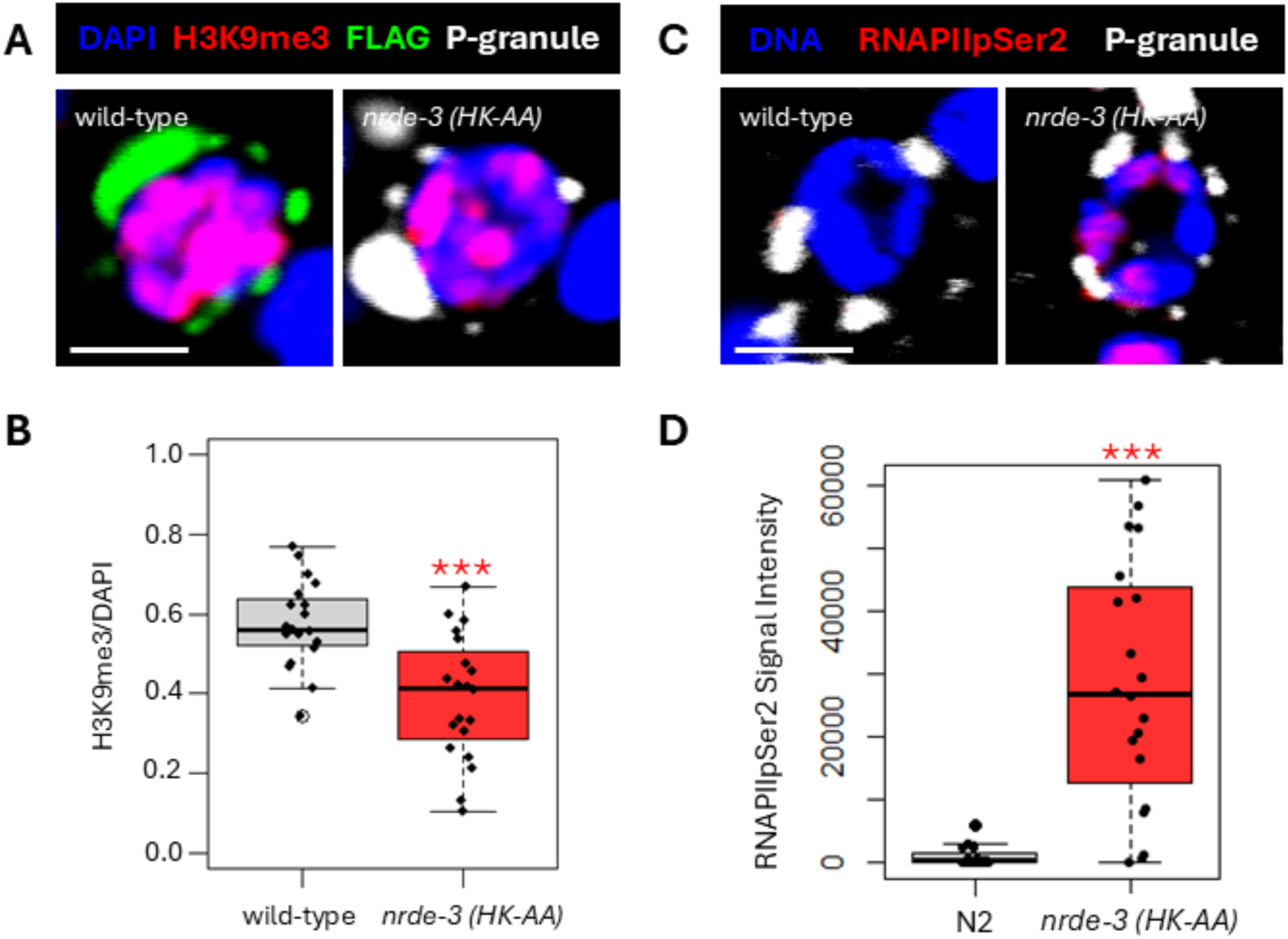
NRDE-3 is required for H3K9me3 hyperdeposition and the absence of functional NRDE-3 allows for aberrant transcription in Z2/Z3 of starved L1s. **(A)** Z2/Z3 of starved L1s were stained for DAPI (blue), P-granules (white), and H3K9me3 (red) in wild-type and *nrde-3 (HK-AA)* mutant animals. Scale bars: 5 μm. **(B)** Quantification of H3K9me3 signal overlap with DAPI for Z2/Z3 of starved L1s for data presented in (A). n=20; ***p≤0.001 **(C)** Z2/Z3 of starved L1s were stained for DAPI (blue), P-granules (white), and RNAPIIpSer2 (red) in wild-type and *nrde-3 (HK-AA)* mutant animals. Scale bars: 5 μm. **(D)** Quantification of normalized raw integrated density for data presented in (C). n=20; ***p≤0.001

### NRDE-3 is required for PGC GTS during L1 starvation

Our finding that Z2/Z3 H3K9me3 hyperdeposition is controlled by the NRDE pathway during L1 starvation suggests that NRDE is also required for PGC GTS. To resolve this, we stained starved L1s, either wild-type or animals expressing *nrde-3 (HK-AA)*, for RNAPIIpSer2. As expected, there was little detectable signal in control samples and, by contrast, *nrde-3 (HK-AA)* mutants showed clearly detectable RNAPIIpSer2 (Fig 6C&D). Based on these data, we conclude that the NRDE pathway is required to globally repress germline transcription when L1s face nutritional stress.

## Discussion

The goals of our study were two-fold. The first goal was to define the timing of key early events during germline ZGA. We found that PIE-1 degradation occurs promptly upon the division of P_4_ to form the Z2/Z3 PGCs, and that RNAPII activation occurs simultaneously. Furthermore, we found that if PIE-1 destruction is delayed, via loss of the ZIF-1 ligase, then RNAPII activation is also delayed, showing that PIE-1 destruction is the event that triggers germline ZGA. We note that others have previously shown that PIE-1 is degraded just after P_4_ division and that this is delayed in *zif-1* mutants (Mello et al., 1996; Checchi & Kelly 2006). However, ours is the first study to examine how PIE-1 destruction impacts the timing of RNAPII activation, and this is critical to determine if destruction alone is sufficient to launch ZGA. Regarding PIE-1 proteolysis, the finding that it is only modestly delayed in *zif-1* mutants shows that there is one or more redundantly acting pathways that also target PIE-1 for destruction. The degron for ZIF-1 within PIE-1 has been defined – it is encoded by the zinc finger 1 (ZF1) domain (Reese et al., 2000). Interestingly, recent work has shown that when ZF1 is tethered to heterologous proteins it will promote their destruction in Z2/Z3 in a ZIF-1 dependent manner (Schwartz et al., 2023). These data show that the redundant pathway targeting PIE-1 employs a degron distinct from ZF1, as otherwise ZF1 fusion proteins would be stabilized in Z2/Z3 after loss of ZIF-1, and they are not. Why would nematodes evolve two totally distinct PIE-1 degradation pathways? We propose that because PIE-1 destruction is so important to germline development that multiple pathways evolved to ensure that germline ZGA occurs with proper timing. An additional pathway, targeting a distinct degron, thus buffers the system against both inactivation of the ZIF-1 pathway and mutational inactivation of the ZF1 degron.

We also determined the timeline of the H3K9me3 hyperdeposition in Z2/Z3, first observed in Belew et al., 2021. Data from this study demonstrate that H3K9me3 hyperdeposition initiates shortly after ZGA and is complete by the end of gastrulation (Fig 3). The P_4_ cell shows a H3K9me3 signal pattern similar to what is observed in both somatic cells and in P blastomeres (Belew et al., 2021). In this study we found that the baseline H3K9me3 pattern present in P_4_ begins to expand in Z2/Z3 shortly after ZGA, and hyperdeposition continues through gastrulation (Fig 3). The slight lag between the onsets of transcription and H3K9me3 hyperdeposition suggests that hyperdeposition is dependent on ongoing transcription, a prediction strengthened by the outcome of the second goal of this study, to identify the pathway promoting H3K9me3 hyperdeposition.

The temporal connection between ZGA and H3K9me3 hyperdeposition immediately invoked the nuclear RNAi pathway, which has demonstrated a role in coupling active transcription to H3K9me3 modification at nearby nucleosomes (Guang et al., 2010; Burkhart et al., 2011; Burton et al., 2011; Mao et al., 2011). Indeed, we found that mutations in multiple components of the nuclear RNAi pathway (*hrde-1* and *nrde-4*) prevents H3K9me3 hyperdeposition in embryonic Z2/Z3 (Fig 4). The literature documents multiple mechanisms for H3K9me3 deposition. One involves the direct activity of two methyltransferases, where MET-2 deposits H3K9me1/2 and SET-25 finishes up by trimethylating H3K9 (Towbin et al., 2012). The second involves the nuclear RNAi pathway, where NRDE proteins direct SET-25 to targets for trimethylation (Padeken et al., 2021). We see in Figs 4 and 5 that when the NRDE pathway is nonfunctional, H3K9me3 is reduced in Z2/Z3, suggesting that the residual H3K9me3 signal we see was deposited by the MET-2/SET-25 pathway. When SET-25 is removed from the picture altogether, however, no H3K9me3 can be seen. Thus, our work supports existing literature that H3K9me3 can be deposited via multiple mechanisms. We also examined starved L1s, where we have previously shown that SET-25 is required for both H3K9me3 hyperdeposition and for GTS (Belew et al., 2021), and found that NRDE-3 is also required for both events in starved L1s (Fig 6).

One interesting aspect of the data presented here are the differential requirements for NRDE pathway components in initiating versus maintaining H3K9me3 hyperdeposition in PGCs. During initiation, loss of either HRDE-1 or NRDE-4 results in a defect in hyperdeposition whereas maintenance requires NRDE-1, –2, and –3, but not NRDE-4 or HRDE-1. The differential requirements for the secondary AGOs can be explained by expression patterns, as detailed above, however why NRDE-4 shows a differential requirement is not obvious and requires continued investigation. Interestingly, recent work has shown that NRDE-3 promotes RNAPII inactivation during oocyte maturation in the adult gonad (Chen & Phillips, 2025). These data, together with the work reported here, thus supply a new function for nuclear RNAi: GTS of germline gene expression within the germline. This contrasts with another documented role for nuclear RNAi, the suppression of germline gene expression in somatic cells. Thus, nuclear RNAi can silence germline genes both outside and within the germline.

Perhaps the most intriguing result from this study is the hand-off between HRDE-1 and NRDE-3 in directing the NRDE pathway in Z2/Z3 at the embryo-to-L1 transition. HRDE-1 and NRDE-3 are both nuclear Argonautes but are traditionally thought to work separately in the germline and soma, respectively (Seroussi et al., 2022). Our study shows that HRDE-1 and NRDE-3 are directing the same pathway in the same cells, but their activity is tied to different life stages. This phenomenon brings up several questions. One, what sort of programming is changing in Z2/Z3 to mark the transition from embryo to L1? Work from our group has shown that Z2/Z3 can be activated after the L1 animal eats, but the HRDE-1 to NRDE-3 hand-off occurs prior to any feeding-related signaling cues. How is the HRDE-1 to NRDE-3 hand-off occurring? Some clues lie in the expression pattern of the two Argonautes. We know that NRDE-3 is expressed at low levels Z2/Z3 during embryogenesis and becomes more apparent in Z2/Z3 in starved L1s (Fig S2). We note here that the NRDE-3 expression observed by us is slightly different than that reported in Seroussi et al., 2023, where we can clearly detect NRDE-3::GFP in Z2/Z3 in starved L1s. HRDE-1 expression comes up robustly in Z2/Z3 during late embryogenesis and remains present in germ cell nuclei throughout development (Seroussi et al., 2023). Thus, the paucity of NRDE-3 and the presence of HRDE-1 in embryonic Z2/Z3 explain why H3K9me3 hyperdeposition depends on HRDE-1 in these cells. Left unexplained, however, is why HRDE-1 becomes dispensable in starved L1s, despite robust expression. Future work is clearly needed to answer this important question.

Last but not least is the question of what genes are nuclear RNAi targeting in Z2/Z3? We propose that both HRDE-1 and NRDE-3 target the same set of genes in Z2/Z3, and that they are the genes activated at ZGA. This would explain the short time lag between ZGA and the onset of H3K9me3 hyperdeposition. Our data also raise the interesting question of why a system evolved to activate germline transcription in the embryo just to shut it down again when embryos hatch into an environment lacking nutrients. Why not simply delay germline ZGA until L1s have found food? *C. elegans* likely live in a boom-or-bust situation vis-à-vis the presence of nutrients in the environment, and it may be that newly hatched L1s spend considerable time looking for food. Importantly, the Z2/Z3 PGCs are the descendants of cells that were never transcriptionally active. As such, we propose that germline ZGA in the embryo provides a window of opportunity for PGCs to replenish factors that might be needed for PGCs to survive potentially long periods of starvation.

In summary, our work shows that nuclear RNAi is required for both H3K9me3 hyperdeposition in embryonic Z2/Z3 and for H3K9me3 maintenance and GTS in starved L1s. Our data thus suggest a novel transcriptional negative-feedback loop, operating at the genome scale, where germline ZGA triggers both gene expression and H3K9me3 hyperdeposition in embryos and nuclear RNAi then feeds back in starved L1s to shut down the very genes that activated it in the embryo. To strengthen this model it will be important to identify the small RNAs that interact with HRDE-1 and NRDE-3, and such experiments are now in progress.

## Materials & Methods

### C. elegans strains

Strains were maintained at 20°C on nematode growth medium (NGM) seeded with *Escherichia coli* strain OP50 or HT115. List of strains used in this study are listed in Table S1. N2 and MT17463 strains were obtained from Caenorhabditis Genetics Center.

### Egg preparation

Gravid adults were washed from plates with M9 minimal medium (22 mM KH_2_PO4, 22 mM Na_2_HPO_4_, 85 mM NaCl, and 2 mM MgSO_4_). Embryos were extracted using a bleach solution (3.675 ml H_2_O, 1.2 NaOCl, and 0.125 10N NaOH) and by switching off between vortexing for 30 seconds and shaking for 1 minute for a total of 5 minutes. Embryos were then washed 3 times with M9 minimal medium.

### RNAi treatment

HT115 cells transformed with an empty pL4440 vector was used as a negative control. *zif-1* RNAi *E. coli* clone was obtained from the Ahringer library and verified by Sanger sequencing. Animals were grown on HT115 food plates for the first 48 hours, then L4 animals were transferred to *zif-1* RNAi plates for 24 hours. *hpl-2* vector was made by transforming the entire *hpl-2* (isoform c) coding sequence into competent HT115 (DE3) cells. Animals were grown on *hpl-2* RNAi plates for the entirety of their life cycle.

Bacteria were streaked on LB-agar plates containing 100 µg/ml carbenicillin and 12.5 µg/ml tetracycline, then incubated at 37°C overnight. Single colonies were picked and grown in 25 ml LB cultures with 100 µg/ml carbenicillin and 12.5 µg/ml tetracycline. 500 µl of the culture was seeded on 60 mm Petri dishes containing 5 mM IPTG.

### Antibodies and dilutions

<u>GFP</u>: Mouse antibody from Sigma-Aldrich (mab3580) was used at a dilution of 1:100. <u>RNAPIIpSer2</u>: Rabbit antibody from Abcam (ab5095) was used at a dilution of 1:100. <u>H3K9me3</u>: Rabbit antibody from Abcam (ab176916) was used at a dilution of 1:1000. <u>P-granule</u>: Mouse antibody from Developmental Studies Hybridoma Bank (K76) was used at a dilution of 1:20. <u>Secondary antibodies</u>: Alexa Fluor conjugated secondary antibodies from Invitrogen were used at a dilution of 1:200.

### Immunofluorescence staining

Embryos were extracted from gravid adults as described above in “Egg preparation”. L1 animals were obtained by egg preparation, then animals were allowed to hatch overnight in M9 minimal medium on a tube rocker. 100 µl of embryos or L1s were spotted onto a poly-L-lysine coated slide. Samples were allowed to settle onto the slide for 10 minutes before coverslips were applied. Slides were put on dry ice for 30 minutes, then permeabilized by freeze-cracking.

#### For GFP staining experiments

After freeze-cracking, slides were transferred to 100% ice-cold (−20°C) methanol for 30 minutes, then blocked with TNB (100 mM Tris-HCl, 200 mM NaCl, 1% BSA) supplemented with 10% normal goat serum (NGS) for 2 hours at room temperature (RT). Primary antibodies were applied at the dilutions described above in TNB supplemented with 10% NGS. Slides were incubated at 4°C overnight. GFP and RNAPIIpSer2 co-staining experiments were done using the GFP staining protocol.

#### For RNAPIIpSer2 staining experiments

After freeze-cracking, slides were transferred to 100% ice-cold (−20°C) methanol for 2 minutes, then fixing solution (0.08 M HEPES pH 6.9, 1.6 mM MgSO_4_, 0.8 mM EGTA, 3.7% formaldehyde, 1X phosphate-buffered saline) for another 30 minutes at RT. Slides were then washed 3 times with TBS-T (TBS with 0.1% Tween-20), then blocked for 30 minutes with TNB. Primary antibodies were applied at the dilutions described above, and slides were incubated at 4°C overnight.

#### For H3K9me3 staining experiments

After freeze-cracking, slides were transferred to 100% ice-cold (−20°C) methanol for 10 seconds, then fixing solution (as described above) for 10 minutes at RT. Slides were washed 3 times with TBS-T, then blocked with TNB supplemented with 10% NGS for 2 hours at RT. Primary antibodies were applied at the dilutions described above, and slides were incubated at 4°C overnight.

After the primary antibody 4°C overnight incubation, slides were washed 5 times with TBS, then incubated with secondary antibodies and Hoechst 33342 dye for 2 hours at RT. Slides were then washed 5 times with TBS. Mounting medium (50% glycerol in PBS) and coverslips were applied and sealed with Cytoseal XYL.

### Immunofluorescent imaging

All slides were imaged using an Olympus Fluoview FV1000 confocal microscope (60x objective) and Fluoview Viewer software.

### Quantification and statistical analysis

All measurements were done in ImageJ.

Normalized raw integrated density was obtained for GFP and RNAPIIpSer2 quantification. A region of interest (ROI) outlining P_4_ or Z2/Z3 was created in ImageJ using the DAPI channel. The raw integrated density (sum of values in ROI) and ROI area were measured. The raw integrated density was then normalized by the area to obtain the “signal intensity”.

H3K9me3 signal overlap with DAPI was calculated using a ROI outlining P_4_ or Z2/Z3 and a second ROI outlining the relevant H3K9me3 signal from said cell. The area of H3K9me3 signal was then divided by the area of the DAPI ROI outline to obtain the fraction of signal overlap. To correct for the variation between replicates, the fraction of signal overlap was normalized by the average control value.

For statistical analysis, the Shaprio-Wilk test and F-test were used to check for normal distribution and variance. Data were then analyzed using Student’s T-Test or Wilcoxon Rank Sum Test depending on whether the data was parametric or not.

## Acknowledgements

We are grateful to Carolyn Phillips, Erik Griffin, and Craig Mello for the kind gifts of worm strains. Some strains were provided by the CGC, which is funded by NIH Office of Research Infrastructure Programs (P40 OD010440). This work was supported by grants R01GM127477 (W.M.M) and T32HD060549 (E.C.) from the National Institutes of Health.

## Works Cited

1. Baugh LR, Hu PJ. 2020. Starvation Responses Throughout the Caenorhabditis elegans Life Cycle. Genetics. 216(4):837–878. doi:10.1534/genetics.120.303565. PMID: 33268389

2. Belew MD, Chien E, Wong M, Michael WM. 2021. A global chromatin compaction pathway that represses germline gene expression during starvation. Journal of Cell Biology. 220(9):e202009197. doi:10.1083/jcb.202009197. PMID: 34128967

3. Belew MD, Chien E, Michael WM. 2023. Characterization of factors that underlie transcriptional silencing in C. elegans oocytes. Swygert S, editor. PLoS Genet. 19(7):e1010831. doi:10.1371/journal.pgen.1010831. PMID: 37478128

4. Belew MD, Chien E, Wong M, Michael WM. 2024 Oct 3. The topoisomerase II/condensin II axis silences transcription during germline specification in Caenorhabditis elegans. Arbeitman M, editor. G3: Genes, Genomes, Genetics.:jkae236. doi:10.1093/g3journal/jkae236. PMID: 34128967

5. Burkhart KB, Guang S, Buckley BA, Wong L, Bochner AF, Kennedy S. 2011. A Pre-mRNA–Associating Factor Links Endogenous siRNAs to Chromatin Regulation. Lee JT, editor. PLoS Genet. 7(8):e1002249. doi:10.1371/journal.pgen.1002249. PMID: 21901112

6. Burton NO, Burkhart KB, Kennedy S. 2011. Nuclear RNAi maintains heritable gene silencing in Caenorhabditis elegans. Proc Natl Acad Sci USA. 108(49):19683–19688. doi:10.1073/pnas.1113310108. PMID: 22106253

7. Castel SE, Martienssen RA. 2013. RNA interference in the nucleus: roles for small RNAs in transcription, epigenetics and beyond. Nat Rev Genet. 14(2):100–112. doi:10.1038/nrg3355. PMID: 23329111

8. Checchi PM, Kelly WG. 2006. emb-4 Is a Conserved Gene Required for Efficient Germline-Specific Chromatin Remodeling During Caenorhabditis elegans Embryogenesis. Genetics. 174(4):1895–1906. doi:10.1534/genetics.106.063701. PMID: 17028322

9. Chen S, Phillips CM. 2025. Nuclear Argonaute protein NRDE-3 switches small RNA partners during embryogenesis to mediate temporal-specific gene regulatory activity. eLife. 13:RP102226. doi:10.7554/eLife.102226.3. PMID: 40080062

10. Chien E, Michael WM. 2023. Transcriptional repression during spermatogenesis in C. elegans requires TOP-2, condensin II, and the MET-2 H3K9 methyltransferase. MicroPubl Biol. 2023. doi:10.17912/micropub.biology.000933. PMID: 37692088

11. DeRenzo C, Reese KJ, Seydoux G. 2003. Exclusion of germ plasm proteins from somatic lineages by cullin-dependent degradation. Nature. 424(6949):685–689. doi:10.1038/nature01887. PMID: 12894212

12. Fukuyama M, Gendreau SB, Derry WB, Rothman JH. 2003. Essential embryonic roles of the CKI-1 cyclin-dependent kinase inhibitor in cell-cycle exit and morphogenesis in C. elegans. Developmental Biology. 260(1):273–286. doi:10.1016/S0012-1606(03)00239-2. PMID: 12885569

13. Guang S, Bochner AF, Burkhart KB, Burton N, Pavelec DM, Kennedy S. 2010. Small regulatory RNAs inhibit RNA polymerase II during the elongation phase of transcription. Nature. 465(7301):1097–1101. doi:10.1038/nature09095. PMID: 18653886

14. Kim H, Ding Y-H, Lu S, Zuo M-Q, Tan W, Conte D, Dong M-Q, Mello CC. 2021. PIE-1 SUMOylation promotes germline fates and piRNA-dependent silencing in C. elegans. eLife. 10:e63300. doi:10.7554/eLife.63300. PMID: 34003111

15. Kim H, Ishidate T, Ghanta KS, Seth M, Conte D, Shirayama M, Mello CC. 2014. A Co-CRISPR Strategy for Efficient Genome Editing in Caenorhabditis elegans. Genetics. 197(4):1069–1080. doi:10.1534/genetics.114.166389. PMID: 24879462

16. Larson BJ, Van MV, Nakayama T, Engebrecht J. 2016. Plasticity in the Meiotic Epigenetic Landscape of Sex Chromosomes in Caenorhabditis Species. Genetics. 203(4):1641–1658. doi:10.1534/genetics.116.191130. PMID: 27280692

17. Mao H, Zhu C, Zong D, Weng C, Yang X, Huang H, Liu D, Feng X, Guang S. 2015. The Nrde Pathway Mediates Small-RNA-Directed Histone H3 Lysine 27 Trimethylation in Caenorhabditis elegans. Current Biology. 25(18):2398–2403. doi:10.1016/j.cub.2015.07.051. PMID: 26365259

18. Mello CC, Schubert C, Draper B, Zhang W, Lobel R, Priess JR. 1996. The PIE-1 protein and germline specification in C. elegans embryos. Nature. 382(6593):710–712. doi:10.1038/382710a0. PMID: 8751440

19. Padeken J, Methot S, Zeller P, Delaney CE, Kalck V, Gasser SM. 2021. Argonaute NRDE-3 and MBT domain protein LIN-61 redundantly recruit an H3K9me3 HMT to prevent embryonic lethality and transposon expression. Genes Dev. 35(1–2):82–101. doi:10.1101/gad.344234.120. PMID: 33303642

20. Palancade B, Bensaude O. 2003. Investigating RNA polymerase II carboxyl-terminal domain (CTD) phosphorylation. European Journal of Biochemistry. 270(19):3859–3870. doi:10.1046/j.1432-1033.2003.03794.x. PMID: 14511368

21. Reese KJ, Dunn MA, Waddle JA, Seydoux G. 2000. Asymmetric Segregation of PIE-1 in C. elegans Is Mediated by Two Complementary Mechanisms that Act through Separate PIE-1 Protein Domains. Molecular Cell. 6(2):445–455. doi:10.1016/S1097-2765(00)00043-5. PMID: 10983990

22. Schaner CE, Deshpande G, Schedl PD, Kelly WG. 2003. A Conserved Chromatin Architecture Marks and Maintains the Restricted Germ Cell Lineage in Worms and Flies. Developmental Cell. 5(5):747–757. doi:10.1016/S1534-5807(03)00327-7. PMID: 14602075

23. Schwartz AZA, Abdu Y, Nance J. 2023. ZIF-1-mediated degradation of zinc finger proteins in the Caenorhabditis elegans germ line. Goldstein B, editor. GENETICS. 225(3):iyad160. doi:10.1093/genetics/iyad160. PMID: 37647858

24. Seroussi U, Li C, Sundby AE, Lee TL, Claycomb JM, Saltzman AL. 2022. Mechanisms of epigenetic regulation by C. elegans nuclear RNA interference pathways. Seminars in Cell & Developmental Biology. 127:142–154. doi:10.1016/j.semcdb.2021.11.018. PMID: 34876343

25. Seroussi U, Lugowski A, Wadi L, Lao RX, Willis AR, Zhao W, Sundby AE, Charlesworth AG, Reinke AW, Claycomb JM. 2023. A comprehensive survey of C. elegans argonaute proteins reveals organism-wide gene regulatory networks and functions. eLife. 12:e83853. doi:10.7554/eLife.83853. PMID: 36790166

26. Seydoux G, Dunn MA. 1997. Transcriptionally repressed germ cells lack a subpopulation of phosphorylated RNA polymerase II in early embryos of Caenorhabditis elegans and Drosophila melanogaster. Development. 124(11):2191–2201. doi:10.1242/dev.124.11.2191. PMID: 9187145

27. Seydoux G, Mello CC, Pettitt J, Wood WB, Priess JR, Fire A. 1996. Repression of gene expression in the embryonic germ lineage of C. elegans. Nature. 382(6593):713–716. doi:10.1038/382713a0. PMID: 8751441

28. Towbin BD, González-Aguilera C, Sack R, Gaidatzis D, Kalck V, Meister P, Askjaer P, Gasser SM. 2012. Step-Wise Methylation of Histone H3K9 Positions Heterochromatin at the Nuclear Periphery. Cell. 150(5):934–947. doi:10.1016/j.cell.2012.06.051. PMID: 22939621

29. Wang JT, Seydoux G. 2013. Germ Cell Specification. In: Schedl T, editor. Germ Cell Development in C. elegans. Vol. 757. New York, NY: Springer New York. (Advances in Experimental Medicine and Biology). p. 17–39. https://link.springer.com/10.1007/978-1-4614-4015-4_2. PMID: 22872473

30. Youngman EM, Claycomb JM. 2014. From early lessons to new frontiers: the worm as a treasure trove of small RNA biology. Front Genet. 5. doi:10.3389/fgene.2014.00416. http://journal.frontiersin.org/article/10.3389/fgene.2014.00416/abstract. PMID: 25505902

