## Supplemental Figures 1 and 2 for "Sequential action of the nuclear Argonautes HRDE-1 and NRDE-3 promotes global transcriptional silencing during starvation in the nascent *C. elegans* germline"

**Supplemental information**

**Figure S1: Z2/Z3 movement before, during, and after gastrulation.**

Top row: Wild-type embryos at various developmental stages were fixed and stained for DAPI (blue) and P-granules (white). Dotted lines indicate embryo outline. Scale bars: 5 μm.

Bottom row: Schematic of Z2/Z3 positioning in 3D space in the embryo. Gray arrows indicate Z2/Z3 movement. A=anterior, P=posterior, D=dorsal, V=ventral.

**Figure S2: NRDE-3 is expressed in P_4_ and Z2/Z3 during embryogenesis and in starved L1s.**

(A) *nrde-3::gfp* embryos were fixed and stained for DNA (blue), P-granules (white), and GFP (green). Scale bars: 5 μm.
(B) Live imaging of *nrde-3::gfp* starved L1s. Scale bar: 5 μm.
(C) Live imaging of *hrde-1::gfp* embryos. Scale bar: 5 μm.

**Table S1: Strains used in this study**

| **Name used in paper** | **Strain name** | **Genotype** |
| --- | --- | --- |
| N2 | N2 | wild-type |
| *gfp::nrde-3* | JMC237 | *nrde-3(tor131[GFP::3xFLAG::nrde-3]) X* |
| *nrde-3 (HK-AA)* | USC1563 | *nrde-3(tor131[GFP::3xFLAG::nrde-3], cmp324[HK-AA]) X* |
| *gfp::hrde-1* | JMC231 | *hrde-1(hzhCR176[gfp::3xflag::hrde-1]) III* |
| *hrde-1* | YY538 | *hrde-1(tm1200) III* |
| *set-25* | MT17463 | *set-25(n5021) III* |
| *nrde-1* | YY160 | *nrde-1(gg88) III* |
| *nrde-2* | YY186 | *nrde-2(gg91) II* |
| *nrde-4* | YY453 | *nrde-4(gg129) IV* |
| *pie-1::gfp* | WM330 | *pie-1(ne4301[pie-1::GFP]) III* |
| *zif-1-/-* | EGD410 | *pie-1(ne4301[pie-1::gfp]); zif-1(egx5) III* |

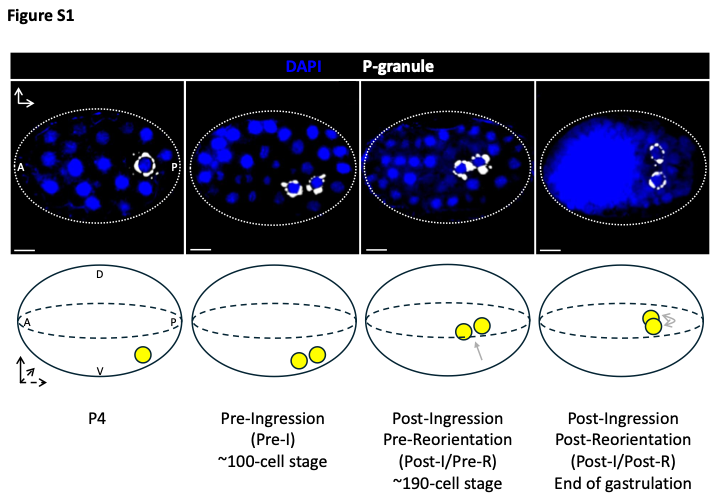

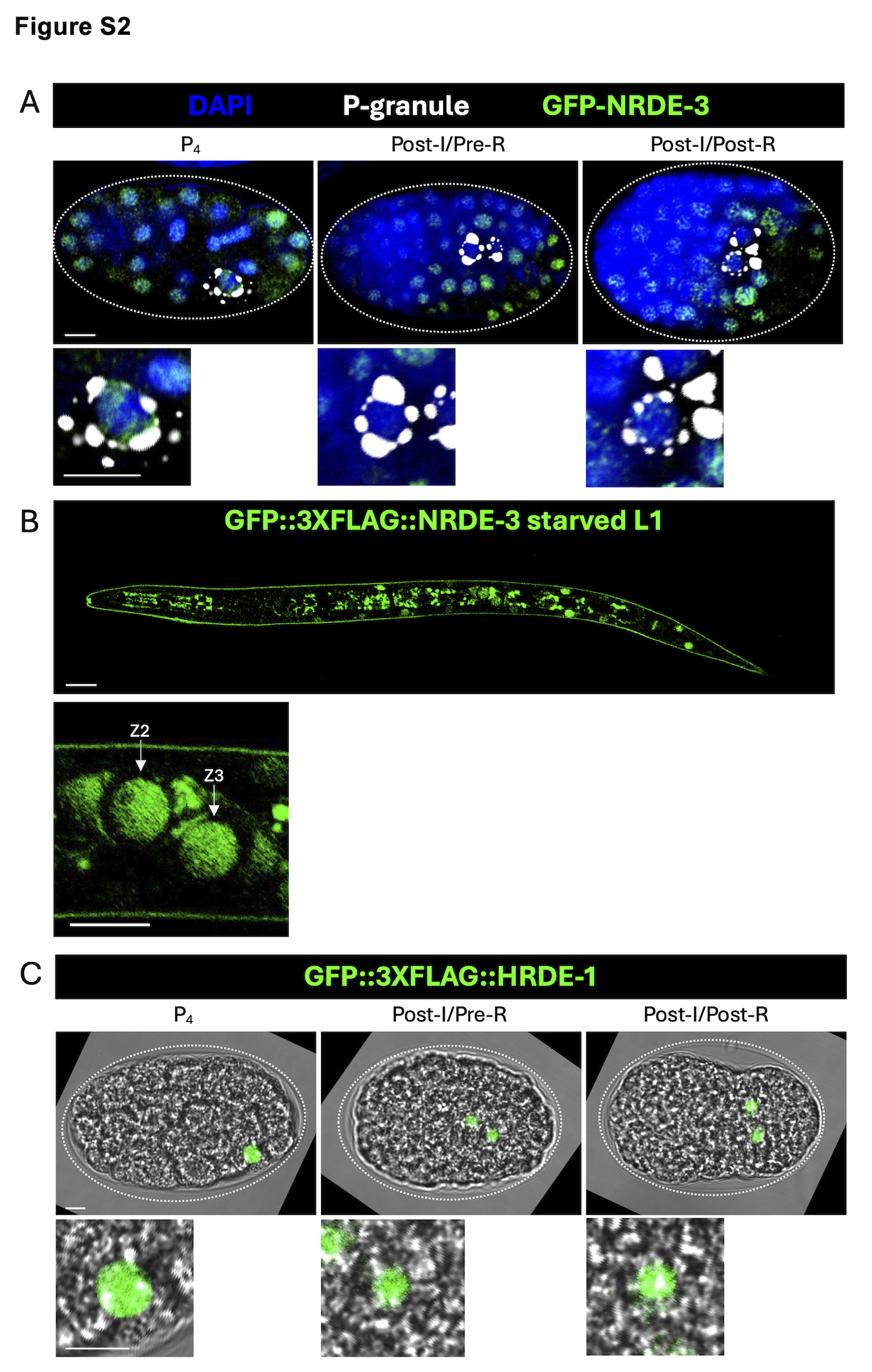
